# Beyond ROIs: Laminar fMRI Mapping with Cylarim

**DOI:** 10.1101/2025.03.25.645237

**Authors:** Gabriele Lohmann, Julius Steiglechner, Lucas Mahler, Klaus Scheffler

## Abstract

Laminar fMRI offers a powerful approach to investigate brain function by resolving neural activity across the distinct layers of the cortical gray matter. However, current laminar fMRI analysis methods often rely on predefined regions of interest (ROIs), which introduce subjectivity and bias due to arbitrary spatial delineation. To address this limitation, we present “Cylarim”, a novel software package that eliminates dependence on rigid ROI definitions. Cylarim employs small, overlapping cylinders that systematically traverse the cortical ribbon, enabling unbiased, large-scale mapping of laminar-specific hemodynamic activity. This approach supports comprehensive sampling of cortical depth-resolved signals across extensive regions. By circumventing the need for manual or atlas-driven ROI selection, Cylarim facilitates the study of feedforward and feedback pathways in diverse cortical areas without compromising spatial coverage. Preliminary results demonstrate Cylarim’s ability to accurately localize layer-dependent responses, providing a robust framework for advancing ultrahigh-resolution fMRI research in neuroscience.

## 1. Introduction

Functional magnetic resonance imaging (fMRI) studies investigating laminar-specific activity have emerged as a crucial tool for understanding the directional flow of information within the brain. By resolving neural activity across the distinct layers of the cortical gray matter, these studies offer insights into the hierarchical processing underlying brain function [1–7], see also the many contributions in [8]. While various methodologies have been proposed for laminar fMRI analysis (e.g. [9–11]), a significant limitation persists: the reliance on user-defined regions of interest (ROIs). This dependency introduces subjectivity and potential bias, as analysis outcomes become highly sensitive to the arbitrary spatial boundaries established during ROI selection. Furthermore, many existing methods restrict their analysis to narrow, relatively straight cortical strips, excluding the valuable information contained within curved cortical regions. To address these challenges, we introduce Cylarim, a novel software package that eliminates the need for predefined ROIs and accommodates the complex geometry of curved cortical surfaces. Cylarim employs an iterative approach, systematically applying small, overlapping cylinders along the cortical ribbon. This cylinder-based strategy, reflected in the software’s name, facilitates the generation of a comprehensive, data-driven map of laminar-specific activity across the entire cortical mantle, enabling a more objective and spatially inclusive analysis of laminar fMRI data. Functional magnetic resonance imaging (fMRI) studies investigating laminar-specific activity have gained significant attention in recent years. These studies hold the potential to elucidate the directionality of information flow between brain regions.

## 2. Cylarim: The Analysis Pipeline

The Cylarim analysis pipeline consists of the following steps:

1. Upsampling to a higher spatial resolution (if needed).
2. Identification of the cortical rims.
3. Calculation of cortical depth normalized to [0,1].
4. Definition of cylinders that traverse the cortical ribbon.
5. Statistical evaluation of laminar activations within individual cylinders.
6. Laminar mapping.

Note that steps 1-3 are identical to the processing pipeline used by the LayNii package [11], while steps 4-6 are novel and specific to Cylarim. In the following, each of these steps will be described in detail. We will illustrate our approach using a test image (“rim M.nii.gz”) which is provided as part of the LayNii package [12].

### Step 1: Upsampling (optional)

The human cerebral cortex exhibits regional variations in thickness, averaging roughly 2.5 mm. While some regions, such as the calcarine cortex, can be as thin as 1 mm, others, like the precentral gyrus and superior frontal and temporal lobes, reach up to 4.5 mm [13]. A significant challenge in functional magnetic resonance imaging (fMRI) is that its typical spatial resolution often falls short of resolving these fine-grained cortical details. To mitigate this, a common practice involves upsampling the fMRI data to a higher spatial resolution, typically around (0.2*mm*)^3^. It is crucial to acknowledge that this upsampling procedure does not generate genuinely higher-resolution information. Instead, it serves as a pragmatic, albeit imperfect, method to improve data visualization and analysis.

### Step 2: Cortical rims

Cylarim assumes the availability of a high-resolution tissue segmentation, specifically delineating grey matter (GM), white matter (WM), and cerebrospinal fluid (CSF). High-resolution segmentation is a prerequisite for accurate differentiation between these tissue types and for precise voxel classification pertaining to the pial and white matter surfaces of the GM. This accurate tissue segmentation is critical for laminar functional magnetic resonance imaging (fMRI) analysis, as it ensures precise anatomical alignment and spatial depth specificity, both of which are essential for the reliable interpretation of functional data across cortical layers. Based on the tissue segmentation, the cortical rims are marked as either the GM/WM border or the GM/CSF border.

### Step 3: Normalized cortical depth

A crucial step in our pipeline involves computing the cortical depth normalized to a range of [0,1]. The output of this step is analogous to the “metric” output generated by LayNii. However, we propose an alternative algorithm which is based on 3D distance transforms. Distance transforms calculate the distance from each voxel in an image to the nearest boundary or feature of interest, effectively creating a map of those distances. Various implementations of distance transforms are available; in this study, we employ the method described in Kiryati et al. [14]. Importantly, our implementation constrains the distance transform paths to remain within the cortical grey matter.

Specifically, we perform two distance transforms (Figures 1):

**Figure 1:**
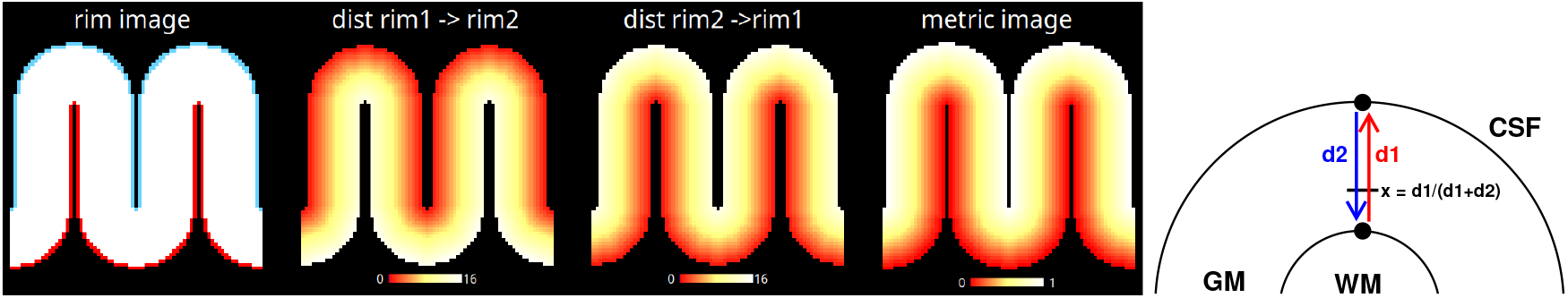
Computing normalized cortical depth using distance transforms. To obtain the normalized cortical depth for every voxel within the cortical grey matter, we perform a constrained 3D distance transform that assigns a distance value for every cortex voxel to the nearest voxel on the border to CSF, and a second distance transform for the reverse distances. Units for the distance transforms are in voxel sizes. The metric image is obtained by combining those two distance images.

- The first, *d*_1_, calculates the distance from the GM/WM border to the nearest GM/CSF border voxel.
- The second, *d*_2_, calculates the reverse distance, i.e. from the GM/CSF border to the nearest GM/WM border voxel.

The normalized metric value, *x*, representing cortical depth, is then computed as:

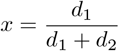

The primary advantage of using distance transforms in this context is their ease of implementation and computational efficiency. Note that this procedure implements an equidistant layering. Below, we will also discuss the equivolume regime.

### Step 4: Cylinders

Cortical cylinders are constructed by pairing corresponding voxels from the grey matter/white matter (GM/WM) and grey matter/cerebrospinal fluid (GM/CSF) borders. For each GM/WM voxel, we identify its nearest neighbor on the GM/CSF surface, and reciprocally. To ensure accurate pairing, this process is performed bidirectionally. As shown in Figure 2, high curvature regions can result in distant nearest neighbors if only one direction is considered. Therefore, the bidirectional approach is essential for complete cortical coverage. Any redundant voxel pairs, identified in both directions, are then removed post-hoc, leaving a set of unique pairs that define the central axes of our cortical cylinders.

**Figure 2:**
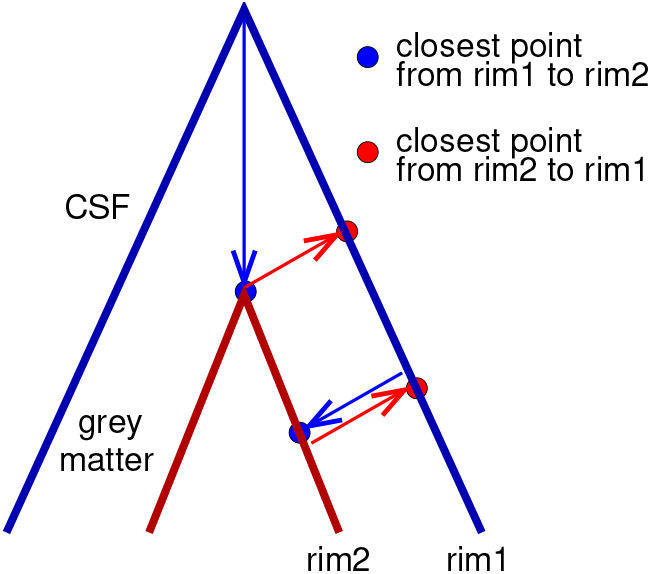
Defining cylinders. Cylinders are defined by connecting corresponding voxels on opposing cortical rims. Due to curvature-induced discrepancies in closest-point mappings between rim1 and rim2, connections are established bidirectionally. For each voxel on rim1, the closest voxel on rim2 is found, and vice versa. The connecting lines are dilated by a user-defined radius, and redundant lines are removed to ensure accurate and complete coverage.

**Figure 3:**
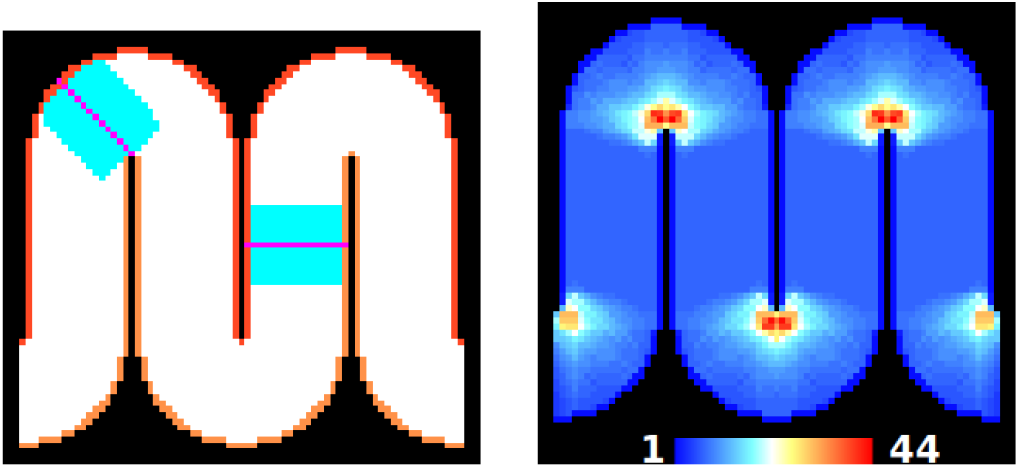
Illustration of cylinders in a synthetic image with a voxel size of (0.1mm)^3^. Two cylinders are shown with their center lines marked in magenta (left). The cylinder radius is 0.7mm. There are 2 × 237 = 474 voxels on the two rims of this image. After excluding duplicate pairs of rim voxels, this yields a total of 343 cylinders for this image, each containing about 200 voxels. The right image shows the degree of overlap of cylinders. The colors encode the number of cylinders by which a voxel is covered. On average, voxels inside the simulated cortical ribbon are covered by about 9 different cylinders, the maximum is 44.

#### Equivolume layering

As highlighted by Waehnert et al. [15] and originally observed by Bok [16], cortical layer thickness varies across the cortical surface, particularly in high-curvature regions where it deviates significantly from an equidistant distribution. To address this, equivolume layering methods have been developed, aiming to ensure each layer contains an equal volume of grey matter [11].

Here we propose a novel equivolume adjustment method based on the idea that at areas of high curvature depth values are not equally distributed within a cylinder. Specifically, for each cylinder, we compute a histogram of the normalized depth values and apply histogram equalization resulting in a uniform distribution of the depth values (Figure 4). Finally, the adjusted depth values from overlapping cylinders and mapped back to the “metric” image.

**Figure 4:**
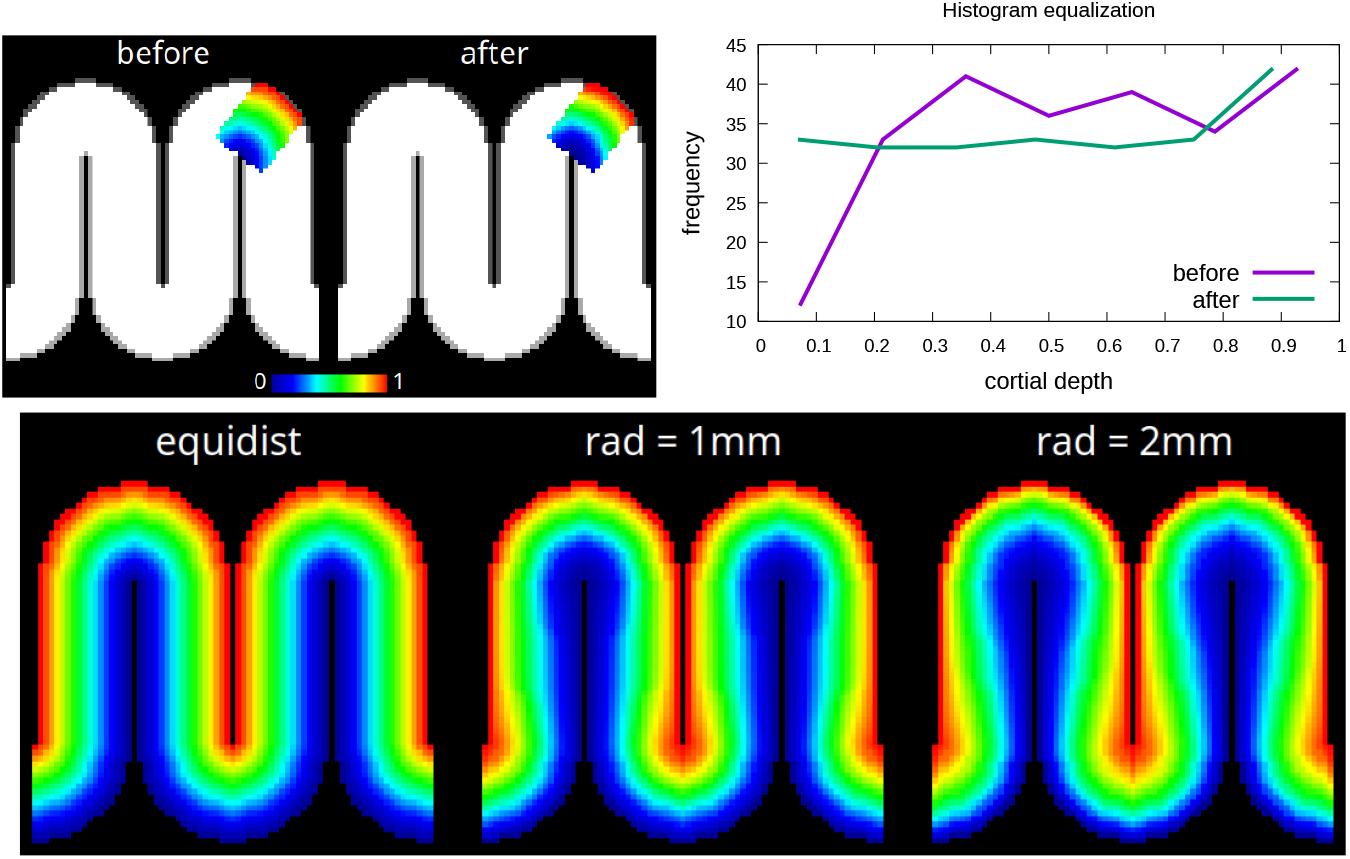
Equivolume adjustment via histogram equalization. The top row shows the effect of histogram equalization (before and after) in a single cylinder. The bottom row shows the effect of equivolume adjustment after averaging all overlapping cylinders. Note that larger cylinder radii lead to a more pronounced equivolume adjustment.

### Step 5: Laminar statistics in a single cylinder

This section focuses on laminar statistics within a single cylindrical volume. Later on, we will detail the methodology for aggregating these single-cylinder results into comprehensive laminar statistical maps. We assume the existence of an fMRI activation map (Zmap) that represents the statistical significance of a specific experimental contrast, illustrated by the z-values associated with, for instance, ‘right-hand fingertapping’ versus ‘left-hand fingertapping’. We propose two alternative methods. The first is based on simple layerwise averages, the second uses a general linear model. To optimize computational speed, the single-cylinder analysis can be performed in parallel across all cylinders.

#### Method 1: Slabs

To analyze laminar differences, we initially divide the cortical ribbon into three equal slabs and calculate the mean z-value within each slab (Figure 5). We then assess the statistical significance of differences between slabs by performing pairwise two-sample t-tests, yielding three contrasts: deep-middle, deep-superficial, and middle-superficial. The statistics obtained for each cylinder using this method are presented in Table 1.

**Table 1:** Laminar statistics calculated in each cylinder. For ease of presentation, these statistics are referred to as μ_1_, μ_2_, μ_3_, and z_1_, z_2_, z_3_. In the slab-based approach, μ_1_, μ_2_, μ_3_ correspond to average fMRI-based activation values within a cylinder. In the GLM-based approach, they are β-estimates.

|  |  |
| --- | --- |
| Mean deep layer ($\mu_1$) | Twosample $t$-test: deep - middle ($z_1$) |
| Mean middle layer ($\mu_2$) | Twosample $t$-test: deep - superficial ($z_2$) |
| Mean superficial layer ($\mu_3$) | Twosample $t$-test: middle - superficial ($z_3$) |

**Figure 5:**
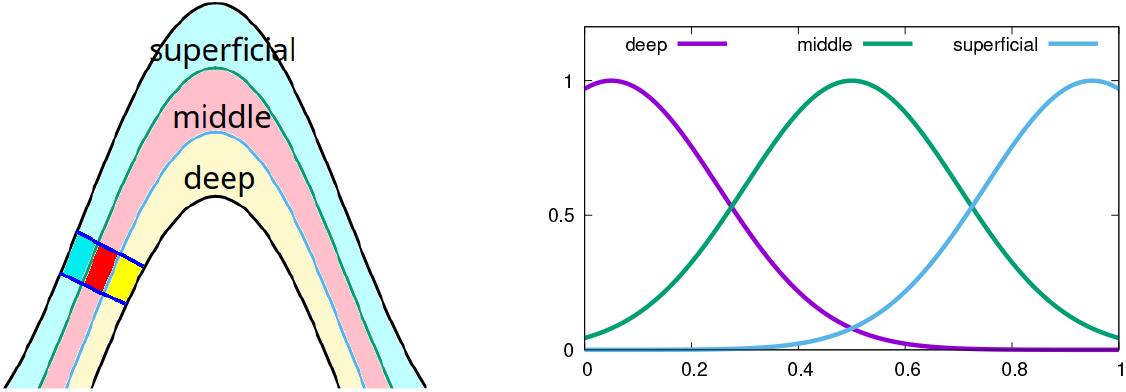
Two approaches for single-cylinder statistics. We propose two alternative methods. The first method is based on three depth-related slabs (left). The means of the fMRI-based z-values are calculated separately in each slab, and pairwise twosample t-tests are performed comparing the means of the three slabs. The cylinder is highlighted by saturated colors. The second method uses a general linear model for each cylinder with three basis functions representing deep, middle and superficial layers (right).

Determining accurate p- and z-values necessitates an estimate of the degrees of freedom, which is typically calculated as the number of independent observations minus the number of covariates. However, the voxels within the Zmap are likely strongly correlated, particularly if the data underwent upsampling. Consequently, the effective number of independent observations is significantly reduced compared to the total number of voxels in the cylinder, making the calculation of reliable p-values for these two-sample t-tests challenging.

Permutation tests offer a solution to the problem of estimating degrees of freedom. By constructing a null distribution that preserves the data’s inherent properties but randomizes the relationship with the test statistic, we can bypass this estimation. A critical requirement for this approach is the exchangeability of observations under the null hypothesis [17, 18].

We use permutation testing to generate a null distribution by randomly shuffling the assignment of voxels to the three depth-related slabs and recalculating the three pairwise two-sample t-tests. The resulting p-values, representing the probability of observing our test statistic under the null hypothesis, are calculated as:

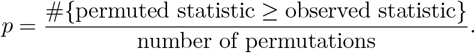

These p-values are then converted to z-scores using the inverse cumulative distribution function of the standard normal distribution. We used 1000 permutations for our experiments.

#### Method 2: General linear model

Cylarim provides an alternative GLM-based approach to laminar statistics, employing three Gaussian basis functions to model cortical depth strata (Figure 5). These functions represent deep (*B*_0_), middle (*B*_1_) and superficial (*B*_2_) cortical layers, respectively. The GLM regression is mathematically expressed as:

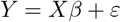

Here, *X* is an *n* × 3 design matrix, where *n* is the number of voxels within the cylinder. The first three columns of *X* encode the basis functions *B*_*j*_, *j* = 0, 1, 2. The observation vector *Y* is mean-centered so that no intercept has to be included into the model. For a given voxel *i* with normalized cortical depth *x*, the corresponding matrix element is defined as *X*_*i,j*_ = *B*_*j*_(*x*). The vector *Y* contains the fMRI activation *z*-values for voxels *i* = 0, …, *n −* 1. The regression coefficients *β* are estimated using the pseudoinverse of *X*.

Based on the GLM results, we calculate three contrasts (1,-1,0), (1,0,-1), and (0,1,-1) that correspond to the three contrasts of the slab-based method described before. As customary, we transform these contrasts into *t*-values using the estimated *β* coefficients and their covariance matrix.

To calculate p-values for the twosample t-tests, we again use permutation tests, specifically by randomly reassigning cortical depth values. This randomization simulates the null hypothesis, which states that cortical depth has no impact on the data. Therefore, if the null hypothesis is true, any observed structure in the fitted model should be equally probable for any random shuffling of cortical depth values.

In each permutation, we recalculate the regression coefficients, *β*, by computing the pseudoinverse of 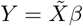. Importantly, the observation vector, *Y*, remains constant across all permutations. Only the design matrix, *X*, is updated to reflect the randomized cortical depth assignments. This ensures exchangeability, allowing us to compute p-values using the same method as for the slab-based approach.

### 2.1 Step 6: Laminar mapping

The statistical values resulting from step 5 are now aggregated through voxel-wise averaging. Specifically, for each voxel, the six statistical measures shown in Table 1 are averaged only across those cylinders that include that voxel. This process yields six distinct statistical maps. This averaging is valid for both mean values and z-scores, which can be assumed to follow a standard normal distribution. Consequently, the output comprises six images, each representing a statistic from Table 1. Note that in the GLM-approach, the first three images contain the averaged *β*-values, instead of the slab-based means.

On the basis of these six images, several additional statistical maps can be generated. For example, to obtain a map that shows cortical areas with a predominant middle layer activation, we compute a conjunction of two contrasts (*−z*_2_ *> z*_1_ & *− z*_2_ *> z*_3_), which corresponds to

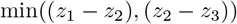

where *z*_1_, *z*_2_, *z*_3_ are as defined in Table 1. Other laminar maps can be obtained in a similar way.

#### Controlling the false discovery rate

We propose to control the false discovery rate (FDR) in the resulting maps by using the Benjamini-Hochberg procedure [19]. This method works by sorting the *p*-values into an ascending order, and naming them *p*_*i*_, *i* = 1, …, *m*, where *m* is the total number of *p*-values. Next, the largest *k* satisfying the condition

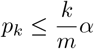

is identified, where *α* is the target FDR level, typically *α* = 0.05. The null hypothesis is rejected for all *i* = 1, …, *k*. This procedure is valid under the assumption of independent or positively correlated tests which aligns with the positive spatial correlation structure commonly observed in fMRI data.

## 3. Experiments

### Simulated data

We applied Cylarim to a simulated Zmap (Figure 6). This Zmap is geometrically aligned with the data set provided by the LayNii-website (“rim M.nii.gz”) that we have used in the previous sections. The simulated Zmap contains activations in the three layers (deep, middle, superficial) as well as combinations of activations across two layers. We have applied permutation tests using 1000 permutations to obtain the three statistics *z*_1_, *z*_2_, *z*_3_ described in Table 1. Here, we have used the slab-based method (Method 1). Figure 6 shows two conjunctions highlighting areas with “deep *>* middle & deep *>* superficial”, as well as “superficial *>* deep & superficial *>* middle”.

**Figure 6:**
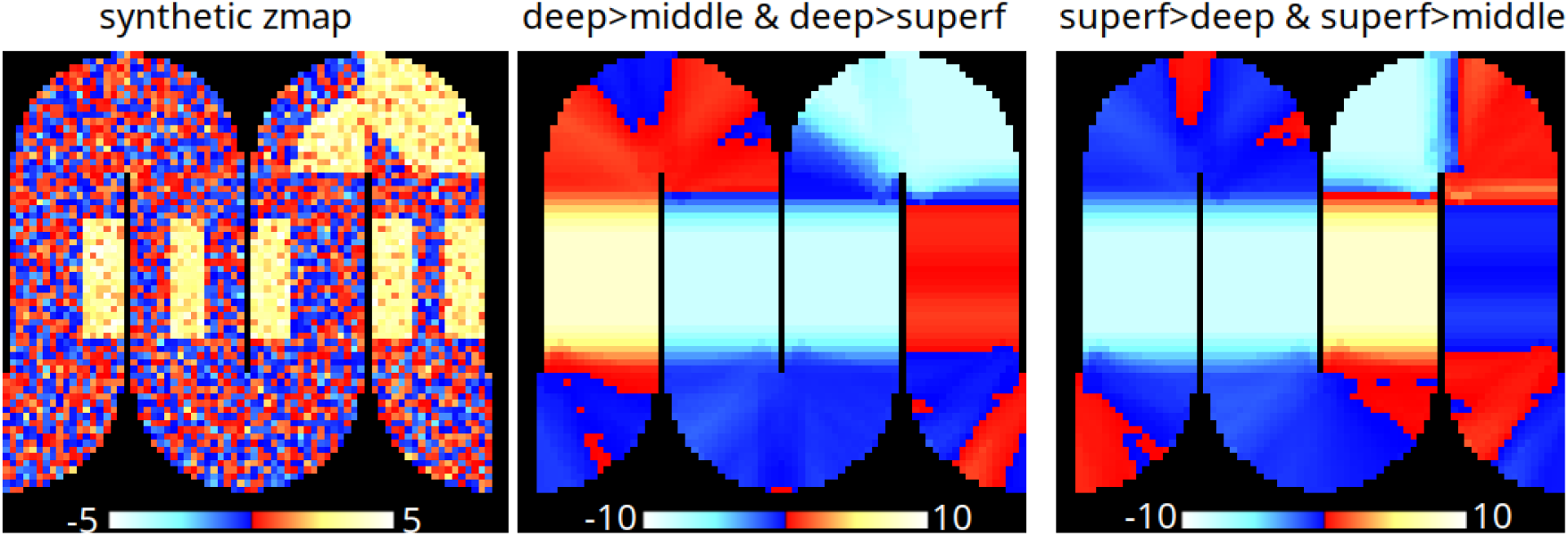
Results using simulated data. The images show the results using the slab-based approach with two conjunctions (deep-middle & deep-superficial), and (superficial-deep & superficial-middle). The results were obtained using 1000 permutations of the slab-based approach (“Method 1”). The colorbars represent z-values uncorrected for multiple comparisons.

### Real data

In addition, we applied Cylarim to another data set provided by LayNII (“sc VASO act.nii.gz”). This data set was part of a study described in Huber et al. [20] in which participants were asked to perform a finger tapping experiment. The data were acquired on a MAGNETOM 7T scanner (Siemens Healthineers, Erlangen, Germany) using the vascular space occupancy method VASO (TI1/TI2/TR = 1100/2600/3000 ms) [21]. The spatial resolution was 0.2 × 0.2 × 0.32 mm with an image matrix of 648 × 648 × 15. For more information about the acquisition protocol and the experimental design, see [20, 22].

Figure 7 shows the results of GLM-based permutation tests (1000 permutations) with three conjunctions (deep-middle & deep-superficial), (middle-deep & middle-superficial) and (superficial-deep & superficial-middle).

**Figure 7:**
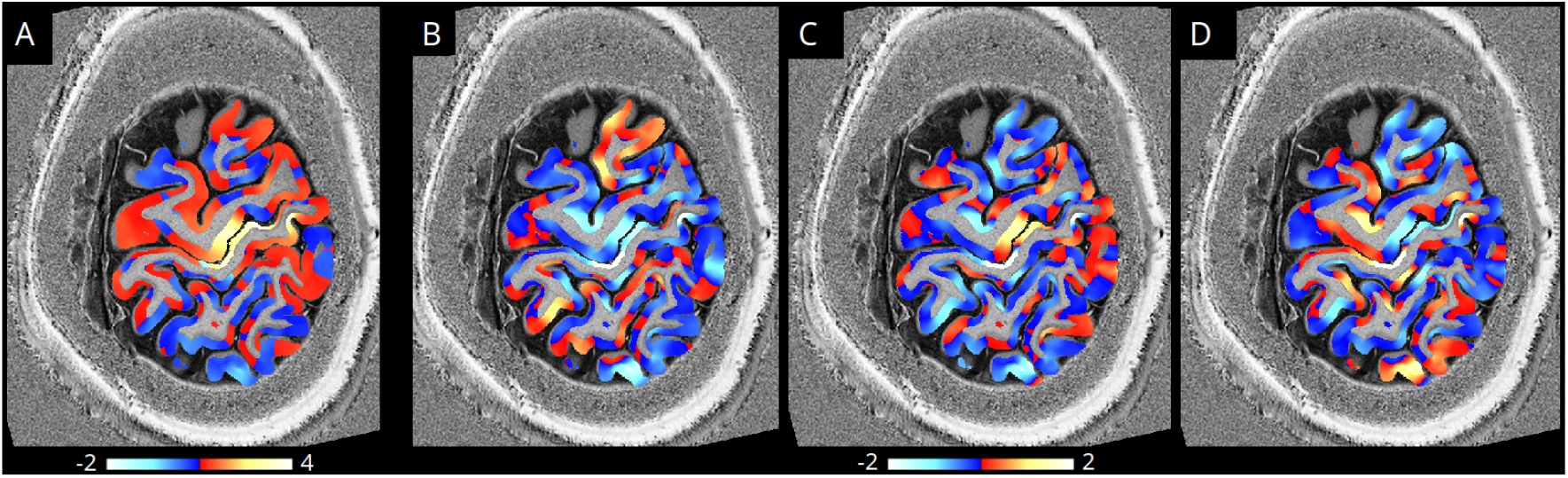
Results (real data) using laminar GLM-based permutation tests. Maximum of the β-estimates indicating areas of a strong laminar-independent contrast in the center of the image (A); Z-values of a conjunction of two contrasts (deep-middle, deep-superficial) (B); Z-values of a conjunction of two contrasts (middle-deep, middle-superficial) (C); Z-values of a conjunction of two contrasts (superficial-deep, superficial-middle) (D);

## 4. Conclusion

In summary, Cylarim offers a novel, ROI-free approach to laminar fMRI analysis, addressing limitations associated with traditional, predefined region-based methods. By employing a systematic, overlapping cylindrical sampling strategy, Cylarim enables unbiased and comprehensive mapping of depth-dependent hemodynamic activity across the cortical ribbon. This method, demonstrated through preliminary results to accurately localize layer-specific responses, provides a powerful tool for investigating cortical microcircuitry.

## Software and Data

The Cylarim software is available for download from: https://lipsia-fmri.github.io The experimental data were downloaded from the LayNii package [12] (Feb 2025).

## Acknowledgements

This work was supported by the following grants: ERC Advanced Grant (No. 834940), and DFG SPP-2041 (DFG LO1728/2-1, SCHE 658/17-1).

